# A decade of disease survey data in a progeny-provenance trial: Dothistroma needle blight in Scots pine

**DOI:** 10.64898/2026.05.12.724484

**Authors:** Annika Perry, Beth Moore, Susan Jones, Sundeep Kaur, Bridget Crampton, Amir Gurung, Jenni Stockan, Joan Cottrell, Joan Beaton, Stephen Cavers

## Abstract

Longitudinal data on disease susceptibility in forest trees are rare but essential for understanding host–pathogen dynamics and genetic variation in susceptibility traits. We present a long-term multisite common garden dataset quantifying susceptibility of Scots pine (*Pinus sylvestris*) to Dothistroma needle blight. The dataset comprises annual disease assessments collected from the same trees across 11 years, spanning 168 families and 21 Scottish provenances. This design enables partitioning of genetic and environmental sources of variation, evaluation of temporal stability in host response, and estimation of variance components and narrow-sense heritability of susceptibility. The data support analyses of phenotypic plasticity, provenance-level responses, and interactions between disease susceptibility and other adaptive traits. This resource will facilitate predictive modelling of host susceptibility under current and future environmental conditions.

## Background & Summary

Understanding and predicting host susceptibility to disease is critically important for the effective management of existing forests and for informing future forest planning under changing environmental conditions. Disease outbreaks can have profound ecological and economic impacts, and the ability to anticipate variation in host response is therefore increasingly central to both conservation and applied forestry. Despite this, progress in understanding tree disease resistance is often constrained by the limited availability and duration of robust, repeated susceptibility measurements. Visual disease assessments, while widely used, can be subjective and are rarely validated against independent molecular estimates of infection, potentially obscuring true patterns of host response.

Longitudinal studies provide a powerful framework for investigating disease dynamics, as they allow temporal variation in host–pathogen interactions to be quantified and enable trait stability to be evaluated across years. Such data are particularly valuable for distinguishing transient responses from consistent phenotypic differences among individuals or genotypes. However, collecting longitudinal data is inherently challenging for large, long-lived organisms such as trees, where disease development, host response, and environmental exposure unfold over extended temporal and spatial scales. Common garden experimental trials are valuable resources for longitudinal studies, enabling repeated measures performed on the same trees. Their design allows genetic and environmental contributions to trait variation to be disentangled, and trials which span multiple sites, families and provenances enable the quantification of phenotypic plasticity alongside genetic components of susceptibility.

Dothistroma needle blight (DNB) is one of the most important diseases of pines (1) caused by the ascomycete fungal pathogen *Dothistroma septosporum* (and occasionally *Dothistroma pini*) and affecting more than 100 species (2) in every continent except Antarctica. Symptoms, including red-brown lesions on needles, can lead to premature needle loss (3,4), growth retardation and, in severe cases, tree death. As pines are major commercial species worldwide, *Dothistroma septosporum* is of significant economic concern and extensive efforts to characterise the pathogen and associated host response have been undertaken in North America (4,5), Europe (6,7), and Australasia (8,9).

Scots pine (*Pinus sylvestris*) is the most widely distributed pine species and is of global ecological and economic importance. There is extensive genetic and phenotypic diversity in the species (6,10–12), unsurprising given the range of environments it occupies. This is coupled with very low population genetic structure and high gene flow across its range, reflecting its high dispersal capability as a wind-pollinated species (13,14). In Scotland, the remnant native populations are typically small, highly fragmented and distributed across an environmentally highly heterogeneous landscape. It has been demonstrated that there is heritable genetic variation in susceptibility to infection by *Dothistroma septosporum* in Scots pine (6) and there is evidence that the pathogen and host have coevolved (7). However, the long-term dynamics and mechanisms underlying the pathosystem are poorly understood, despite their critical importance for evaluating the vulnerability of native and planted forests.

The dataset presented here is a valuable resource to address that knowledge gap. Its longitudinal structure and experimental design allow: 1) evaluation of the drivers of disease severity; 2) identification of genetic variation in responses among provenances; 3) assessment of the temporal stability of disease resistance; 4) estimation of genetic variance components and narrow-sense heritability associated with susceptibility; 5) exploration of interactions with other key adaptive traits such as growth and reproduction; 6) development of predictive models for host susceptibility; and 7) investigation of the mechanisms underlying host responses to disease.

## Methods

### Experimental field trials

Seedling source, nursery environments and field sites are detailed by (15).

The field site in the south of Scotland (FS, latitude 55.603625°, longitude −2.893025°) was visually surveyed annually from 2015 to 2025 (inclusive) except for 2019 and 2020. Replicate field sites in the east (FE, latitude 56.893567°, longitude −2.535736°) and west of Scotland (FW, latitude 57.775714, longitude −5.597181) were surveyed in 2016 and 2022.

### Annual disease surveys

Surveys to record susceptibility to DNB were performed annually in late summer (between 22nd August and 22nd September). Individual trees were visually assessed for the percentage of needles with symptoms of DNB (‘DNB’, symptomatic needles) by applying a scale based on 5 % increments (where 1 % is equivalent to negligible symptoms). It has been observed that trees with high levels of infection lose their needles sooner than those with low levels of infection (3,4), potentially confounding estimates of the proportion of symptomatic needles (very low levels of infection may be recorded where a very susceptible tree has shed all infected needles). For this reason, needle retention (‘NR’) was also recorded at each survey. As annual growth stages are visible in pines, NR could be estimated in half-year increments (in 2016 it was recorded in quarter-year increments). From 2015 to 2017 a single estimate for each of NR and DNB was recorded for each tree annually. As trees grew larger, from 2018 onwards mean values for NR and DNB were obtained from separate estimates for the top, middle and bottom thirds of the canopy of each tree.

Data on symptomatic needles (trait ‘DNB’) were highly left-skewed, so were log(DNB+1) transformed. Where multiple values were obtained per tree (from different crown positions), these were averaged prior to log-transformation. We have termed this trait ‘*direct susceptibility*’ as it directly captures disease symptoms on the host tree.

Needle retention values (trait ‘NR’) were inverted (1-NR) to provide a measure of needle loss: this also ensured that both susceptibility traits were easily and intuitively interpretable (i.e. that low values denote low susceptibility and high values denote high susceptibility). Needle retention, which we assume arises as a cumulative response to infection and may also reflect host response to other needle diseases or a lack of needle drop as a response, we have termed ‘*indirect susceptibility*’.

### Long-term quantification of susceptibility

Direct and indirect long-term disease susceptibility was quantified for each tree at FS using the Area Under the Disease Progress Stairs (AUDPS; (16)), which summarises repeated measurements over time into a single value (‘*direct-LT susceptibility’*, ‘*indirect-LT susceptibility*’). Trees that died over the period 2015 to 2025 were removed from the analysis. For each trait, AUDPS was calculated using the agricolae package (17) in R (18).

For the field sites in which survey data were only collected in 2016 and 2022 (FE and FW), ‘*direct-LT susceptibility*’ and ‘*indirect-LT susceptibility*’ were represented by the mean value for each trait across the two surveyed years. Where data were missing for one of the years, long-term susceptibility was not estimated.

Finally, direct and indirect susceptibility traits were normalised, to a minimum value of 0 and a maximum value of 1, within each site.

### Grouping long-term susceptibility traits

To ensure that these data are also compatible with analyses that require categorical rather than quantitative trait data (for example, when targeting trees to subsample from the extreme ends of trait distribution), we also outline an approach to categorise the long-term susceptibility data into three distinct groups: ‘*low susceptibility*’; ‘*moderate susceptibility*’ and ‘*high susceptibility’* for each trait based on their rank.

The approach assumes that the tails of the distributions reflect the extremes of host response. Susceptibility traits were discretised using percentile ranks calculated across the full distribution of observations at each site. For each tree, a percentile rank was computed based on its position within the ordered trait distribution. Individuals falling within the lower and upper 20 % of the distribution were classified as low susceptibility and high susceptibility, respectively, while all remaining individuals were assigned to the moderate susceptibility category.

### Validation of susceptibility traits using molecular data

Our visual surveys made the assumption that the symptoms observed were predominantly due to the pathogen *D. septosporum*. To validate this, we used a molecular metabarcoding approach at the FS site at timepoint T3 (needles collected 27-28th September 2023). For a full description of molecular methods see (19). Briefly, needle samples were collected for amplicon sequencing on 27-28 September 2023 (the disease survey in that year was conducted on 21-22 September). A subset of trees (N = 200) was selected to represent the full suite of DNB susceptibility responses while retaining the experimental design of the full trial (described in (15)). Uninfected needles (only those that were visually inspected as green and healthy with no apparent lesions or discolouration) were collected from four separate branches around each tree’s crown and sealed within a labelled polythene bag. Needles were lyophilised, homogenised and total cellular DNA was extracted using a modified Qiagen DNeasy 96 kit (10 µl proteinase K was added during the incubation stage). An optimised Illumina metagenomic library preparation protocol was used to amplify the fungal ITS2 region which was sequenced on a NextSeq2000 using a 2 × 300 bp paired end strategy. Raw reads were processed to OTU tables using a qiime2 pipeline (20). Briefly, reads were filtered with fastp (21) to remove short reads (< 90 bp), polyG tails, flanking regions surrounding the target sequence, adapters and reads with high error rates. Remaining reads were denoised with dada2 (22), and classified with classify-sklearn against the UNITE v9.0 database (23) with a 97.5% SH.

The assignment of taxa to the OTUs was verified using massBLASTer against the UNITE and INSD databases. Relative abundances of OTUs per tree, where the number of reads per OTU was presented as a proportion of the total number of reads for each tree, were successfully obtained for all but three of the 200 trees (missing tree ids due to poor quality amplification and sequence data: 7291, 7338, 7474). Where multiple OTUs were assigned to a single taxon, total relative abundance was calculated by summing the relative abundance of all assigned OTUs. Additionally, all *Dothistroma* OTUs were binned as *D. septosporum*, as the closely related pathogen, *Dothistroma pini*, has not been recorded in the UK. This exercise was completed for all needle pathogens that are known to cause needle loss (*Coleosporium tussilaginis; Cyclaneusma minus; Gremmenia infestans; Gremmeniella abietina; Lophodermella conjuncta; Lophodermium conigenum; Lophodermium seditiosum; Pachyramichloridium pini; Rhizosphaera kalkhoffii; Sirococcus piceicola*. See Data Availability section for links to data) to validate that symptoms in the visual survey dataset were due to infection by *D. septosporum* and not to other needle pathogens.

### Data Records

Data are deposited with the Environmental Information Data Centre (https://eidc.ac.uk, 24). Traits measured are detailed in Table 1.

**Table 1.**
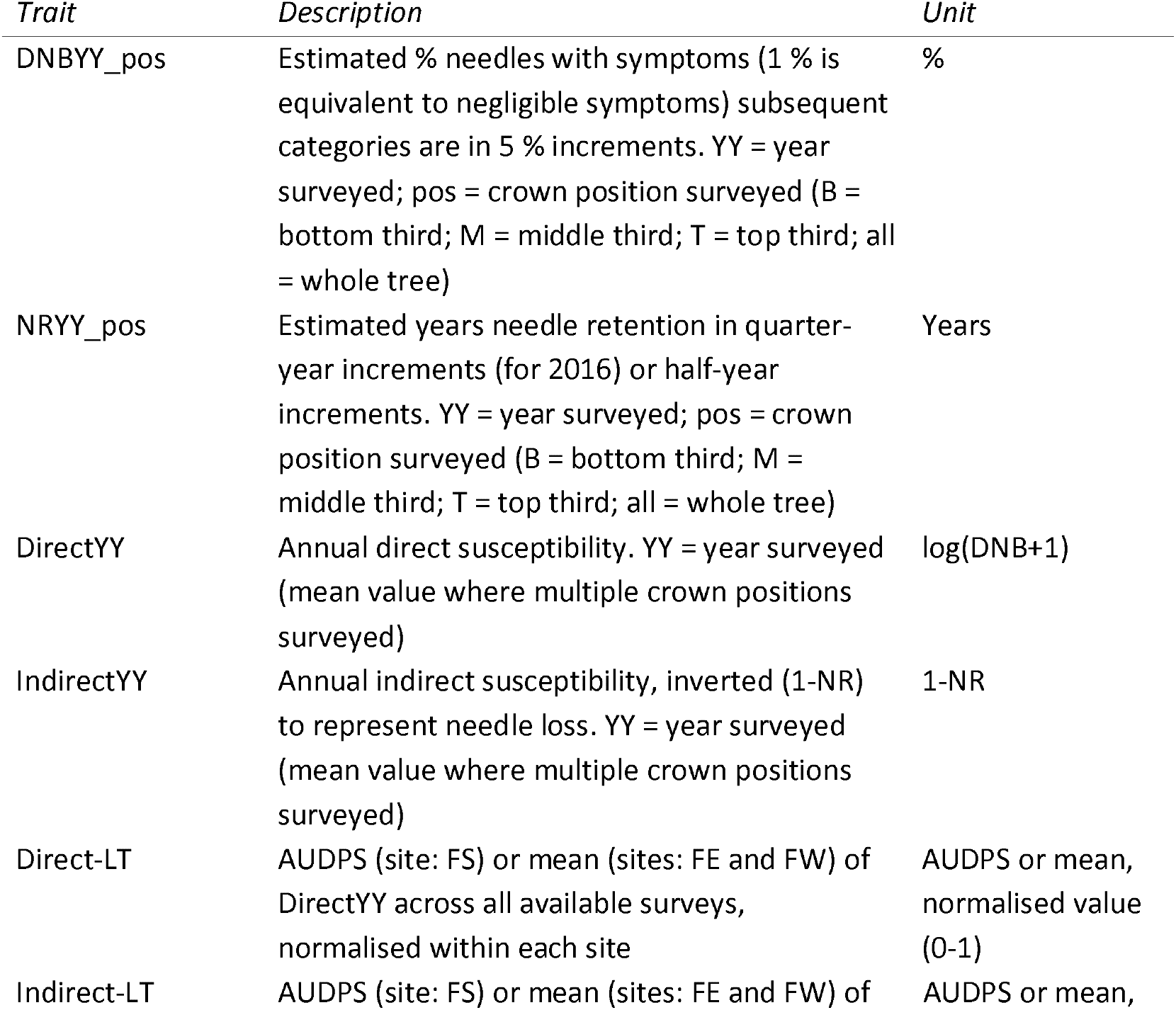
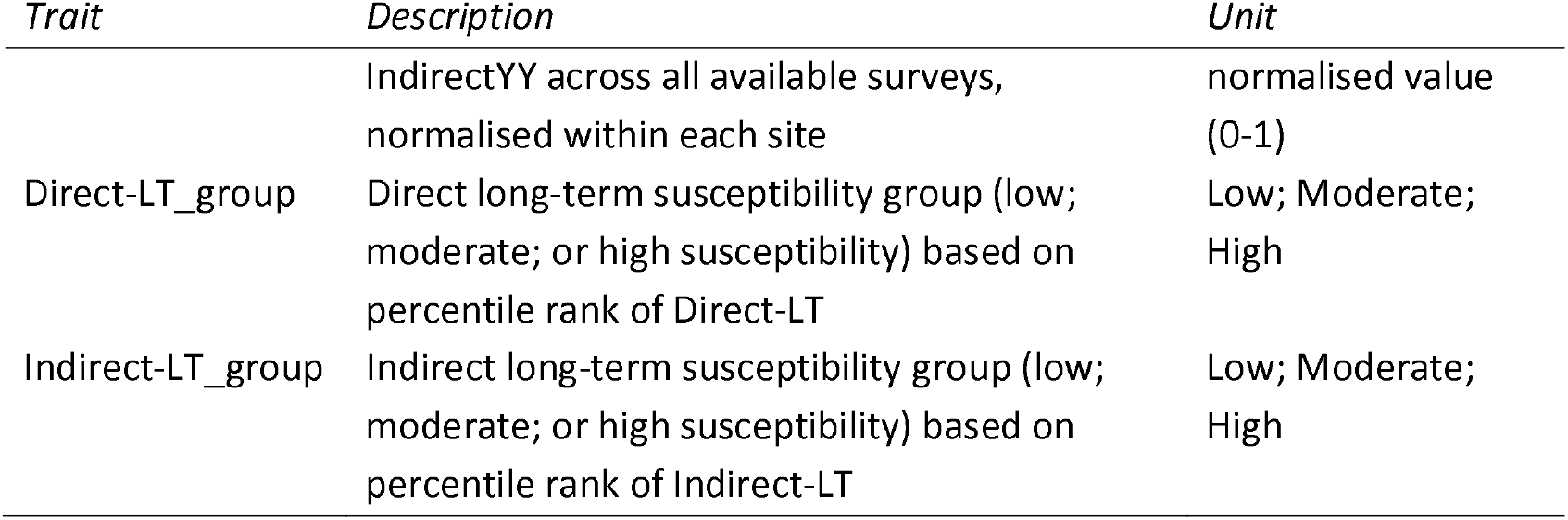
Susceptibility to DNB traits assessed in Scots pine trees at three common garden field trial sites. Where an element of the trait names is in italics, it indicates multiple available options. ‘YY’ = Year, where e.g. 15 is 2015. ‘pos’ = which third of the tree was surveyed: T, top; M, middle; B, bottom; all, whole tree surveyed. Long-term susceptibility traits have the suffix ‘-LT’.

For the dataset, the columns are:

1. PopulationCodeOriginal: Population code originally assigned to the tree
2. FamilyCodeOriginal: Family code (individuals with a shared family code are from the same mother tree) originally assigned to the tree
3. PopulationCodeNew: Population code assigned following genotype checks
4. FamilyCodeNew: Family code (individuals with a shared family code are from the same mother tree) assigned following genotype checks
5. FieldSite: Field trial site where the tree is planted (FE: field site in the east of Scotland, Glensaugh; FS: field site in the south of Scotland, Yair; FW: field site in the west of Scotland, Inverewe)
6. FieldCode: the unique four-digit code identifying each tree links this DNB survey dataset directly with other published datasets (25,26); FE: 5001 to 5672; FW: 6001 to 6004; FS: 7001 to 7672)
7. Block: Randomised block code (A-D)
8. Row: Row that the tree was planted in (numerical)
9. Column: Column that the tree was planted in (numerical)
10. Susceptibility traits: see Table 1 for list of traits and format of column header. Missing data = NA. Data not collected = NC.

### Technical validation

Surveys were performed annually at the same time of year to ensure consistency of the method. Data were checked after each survey and clear errors were immediately corrected by revisiting the tree and re-surveying the trait in question.

We used boxplots to visualise data range and data distribution for each annual susceptibility trait (direct susceptibility and indirect susceptibility) at each field site (Figure 1).

**Figure 1.**
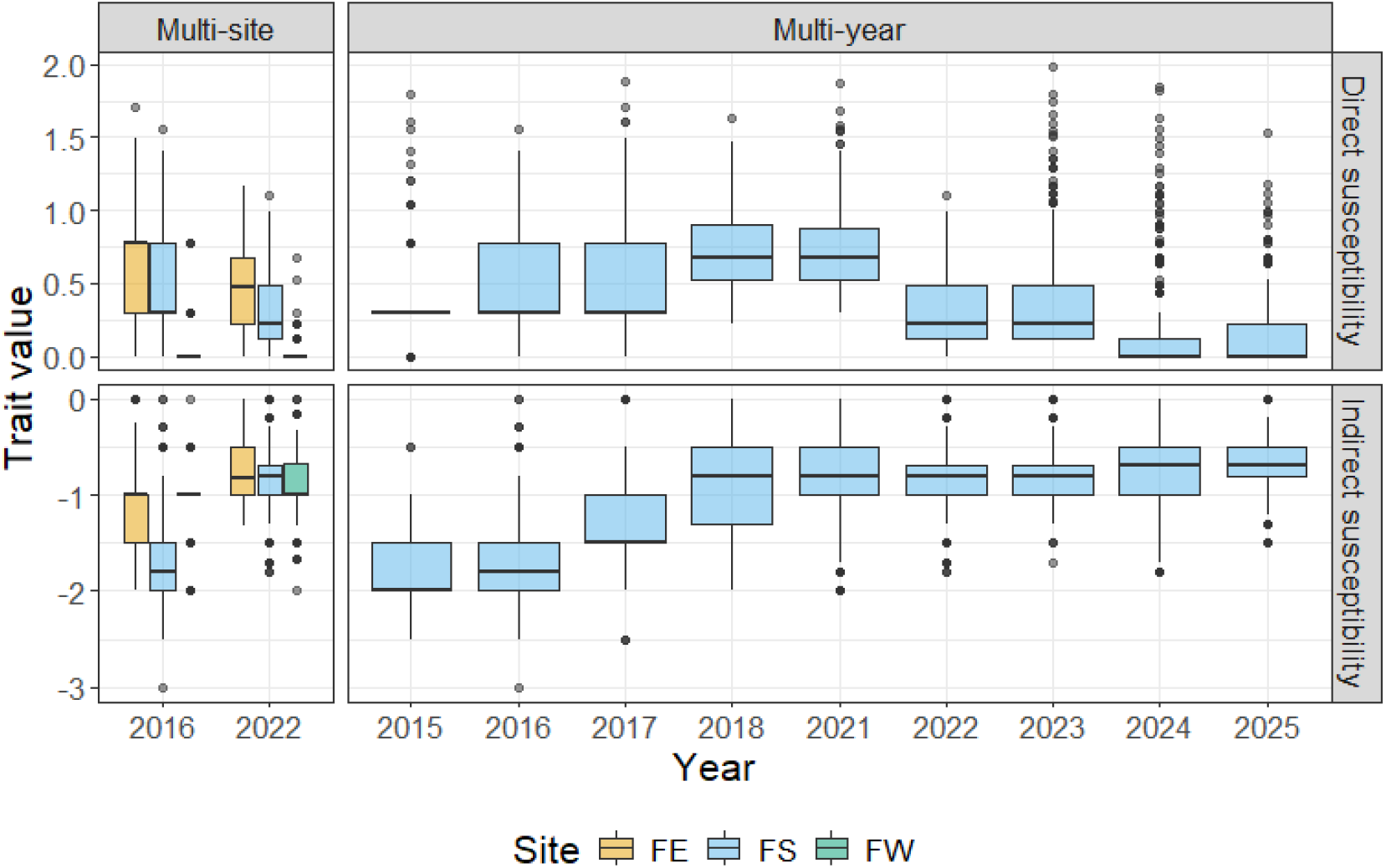
Boxplots showing variation recorded during annual DNB surveys for direct (top panels) and indirect (bottom panels) susceptibility traits. Left: variation among 3 sites in multi-site surveys conducted in 2016 and 2022. Right: variation among years in surveys conducted over the period 2015 to 2025 (excluding 2019 and 2020) at FS. Solid black lines indicate the median trait value. The bottom and top of boxes indicate the first and third quartile. The upper and lower whiskers extend to the highest and lowest values within 1.5 times the interquartile range. Individual points indicate outliers.

We used a pairwise correlation plot to visualise the relationship among normalised longterm susceptibility traits (direct-LT susceptibility and indirect-LT susceptibility) at each field site (Figure 2). The two traits are positively correlated within one another, but the strength of the relationship differs among the sites.

**Figure 2.**
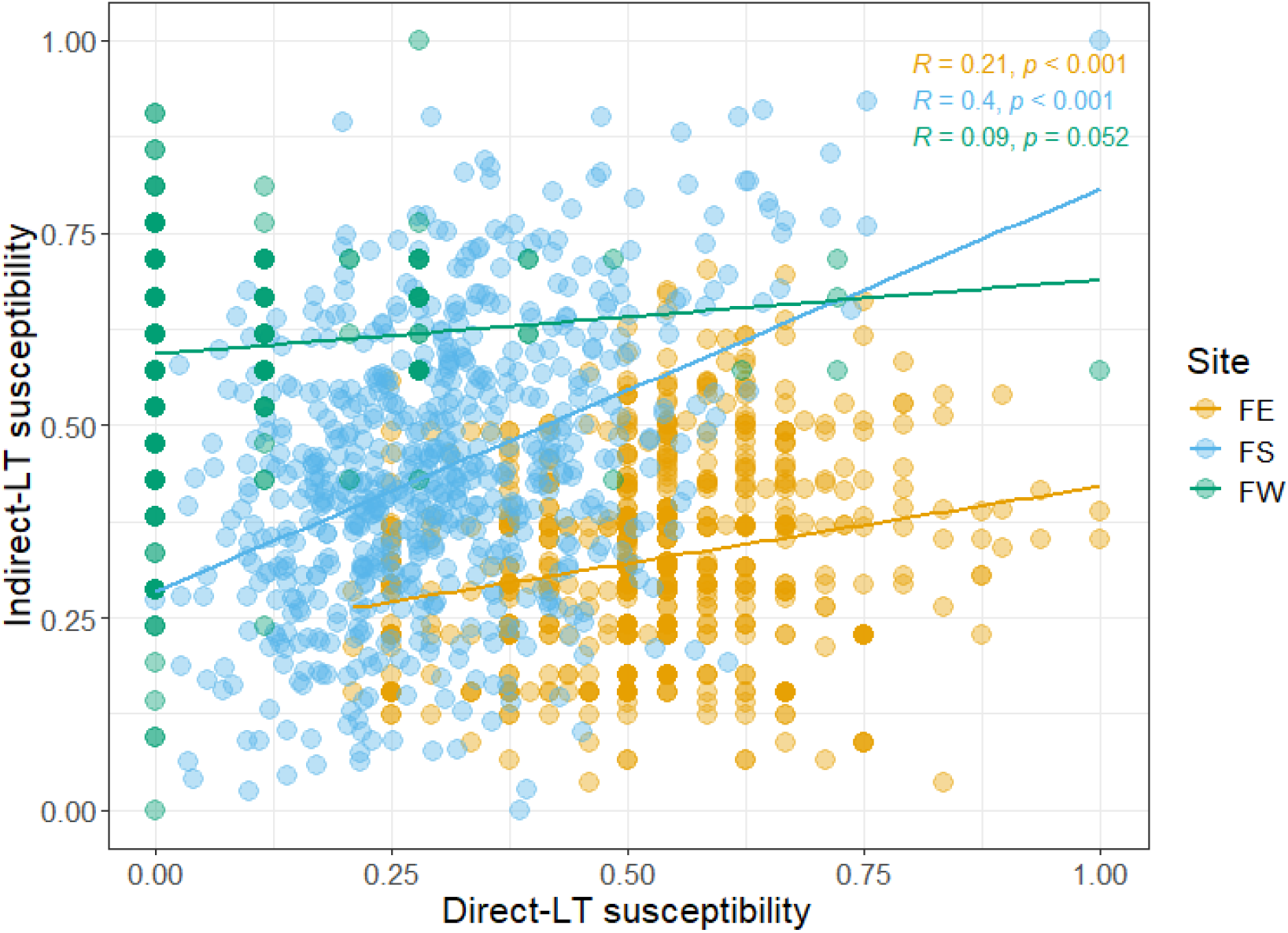
Pearson’s correlation for long-term susceptibility traits (direct-LT susceptibility; indirect-LT susceptibility) quantified for the period 2015-2025 at the FS field site and for 2016 and 2022 at the FE and FW field sites. The correlation coefficient and associated p value are shown for each site.

Discretisation of the long-term susceptibility trait data was applied to allow future analyses that require categorical variables (Figure 3).

**Figure 3.**
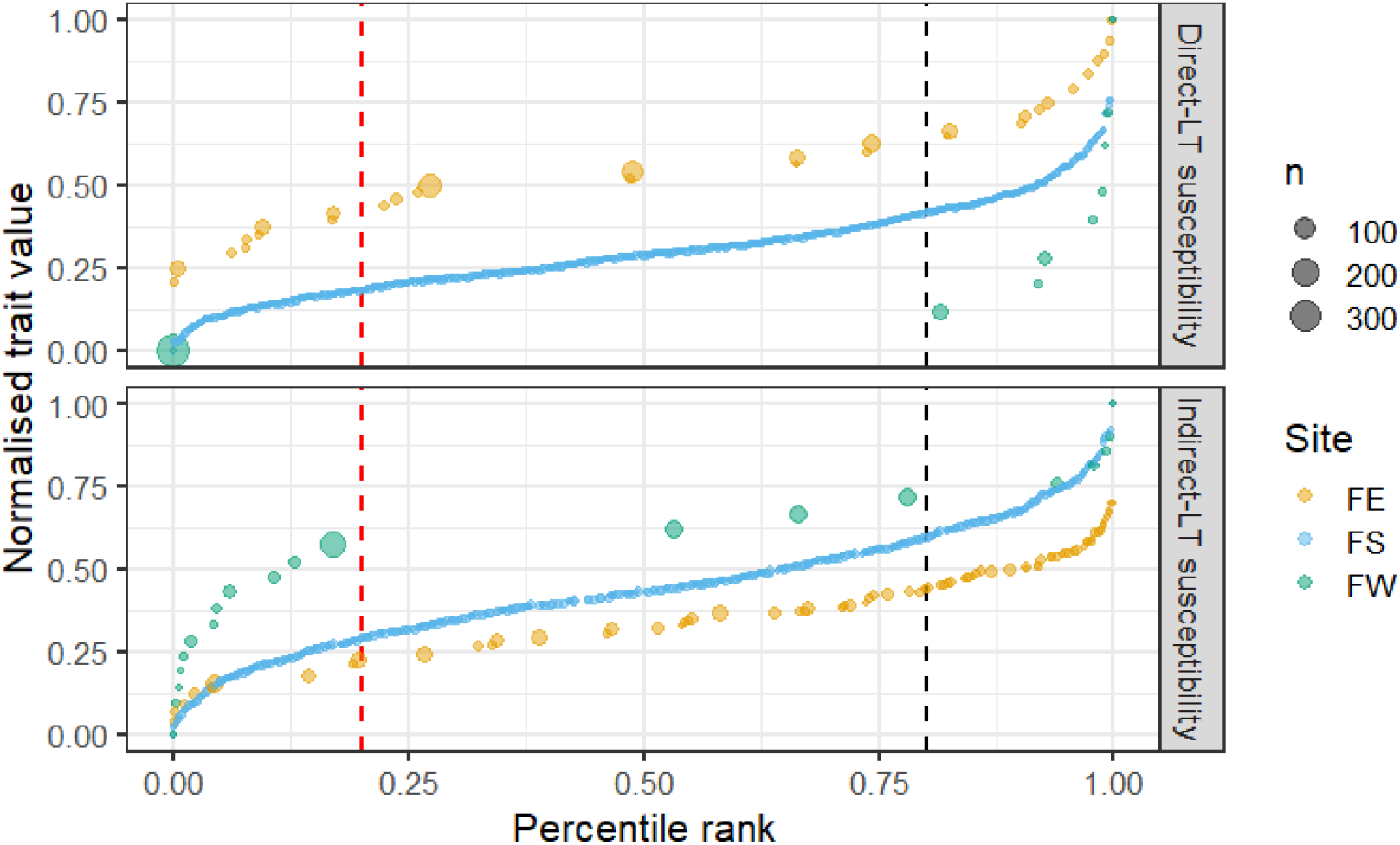
Discretisation of long-term susceptibility data for each site using percentile ranks. The size of the points is relative to the number of trees (n) represented by each trait value. Vertical dashed lines indicate proposed thresholds to group trees into low susceptibility (trees to the left of the red vertical line) and high susceptibility (trees to the right of the black vertical line) with remaining trees in the centre of the distribution assigned to the moderate susceptibility group.

This approach is not recommended for direct-LT and indirect-LT susceptibility traits for trees growing at FW and FE given the skewed and/or patchy distribution of the traits at these sites (Figure 2, Figure 3).

Molecular validation was performed by regressing annual direct and indirect susceptibility traits against the proportion of reads that were identified as *Dothistroma spp*. (Figure 4). This demonstrated that direct susceptibility was highly significantly positively correlated with the relative abundance of in planta *Dothistroma spp*. when the survey is performed in the same year (2023, Figure 3). For surveys performed the year before (2022) or the year after (2024) there were no significant associations between trees’ relative abundance of *Dothistroma spp*. and their direct susceptibility. In contrast, relative abundance of *Dothistroma spp*. was significantly positively associated with indirect susceptibility in all years compared. We therefore consider the direct susceptibility trait to be a robust evaluation of DNB infection in a given year, but not suitable to infer susceptibility in preceding or subsequent years. In these cases, indirect susceptibility and/or long-term susceptibility traits may be preferable.

**Figure 4.**
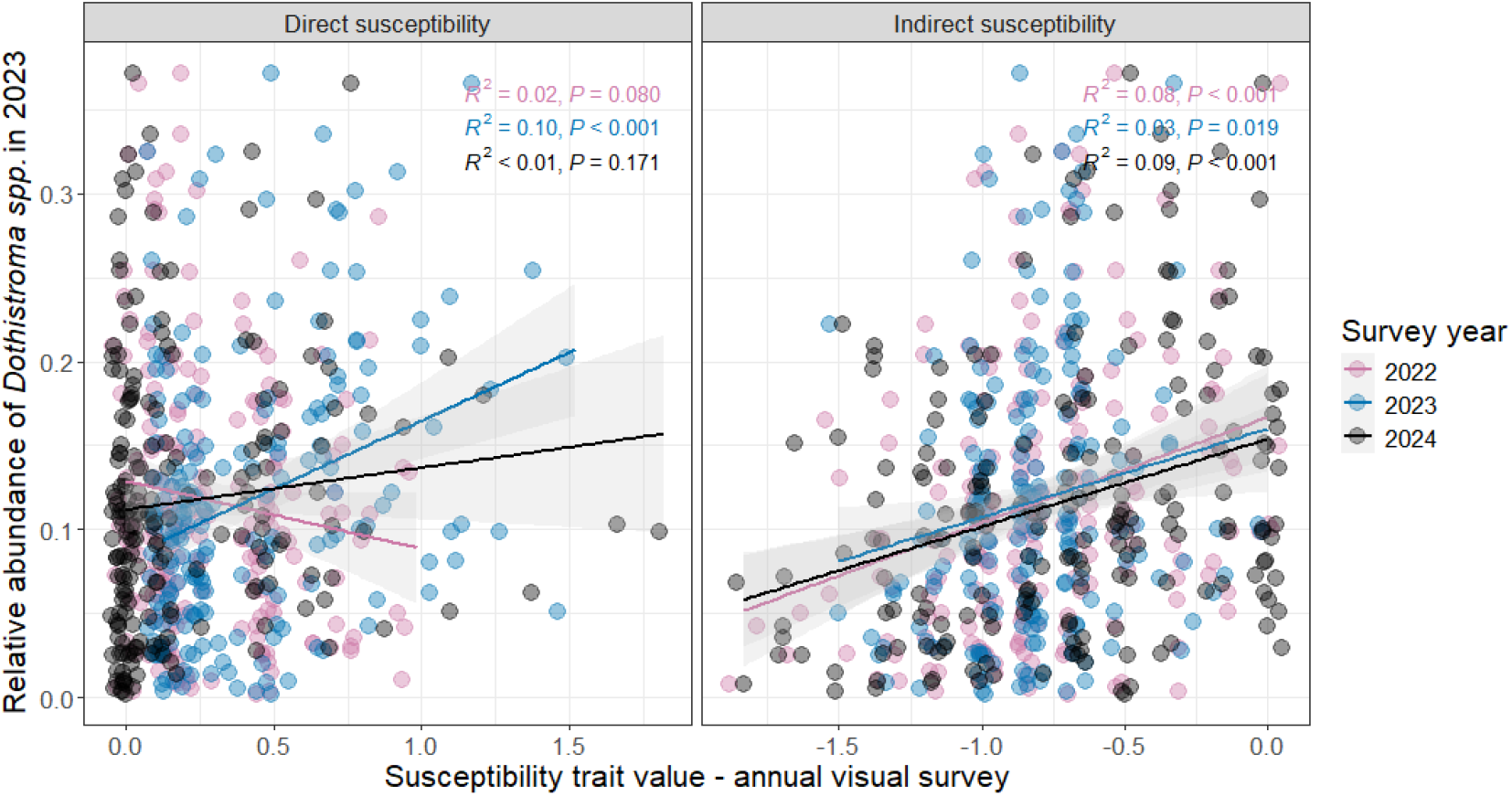
Linear regressions for in planta relative abundance of Dothistroma spp. against visual survey susceptibility traits (direct and indirect susceptibility). Relative abundance of Dothistroma spp. characterised in 2023. Molecular validation was performed using visual survey susceptibility trait data from the preceding year (2022), the same year (2023) and the following year (2024).

Finally, we considered the relative abundance of other needle pathogens relative to those assigned to *Dothistroma spp*. to account for the possibility that observed reported symptoms resulted from host susceptibility to other pathogens (Figure 5). The low relative abundance of other needle pathogens supports the attribution of visual survey of host response to *Dothistroma spp*.

**Figure 5.**
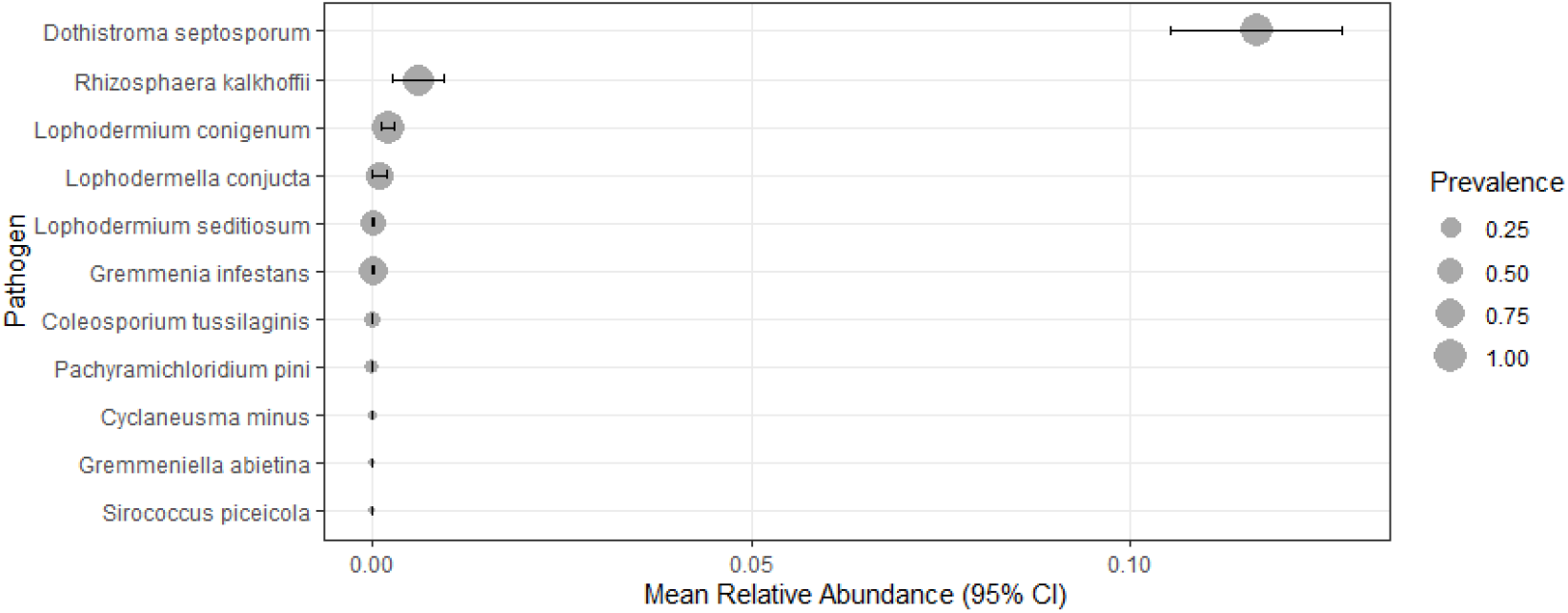
Comparison of relative abundances of needle pathogens in a subset of trees that underwent metabarcoding from FS trial. Bubble position indicates the mean relative abundance, bubble size represents the prevalence of each needle pathogen taxon among all tested trees (N = 197) and the error bars represent the 95% confidence interval.

## Data Availability

The raw metabarcode sequences were deposited into the European Nucleotide archive: https://www.ebi.ac.uk/ena/browser/view/PRJEB88228. The primary filtered OTU table calculated from the ITS2 metabarcoding sequences for T3 are available on GitHub (https://github.com/HuttonICS/PineBiomeDataPaper). Fasta sequences for OTUs assigned to needle pathogen genera and proportion of needle pathogens per tree are available on GitHub https://github.com/HuttonICS/DNB_Survey_Paper_OTU_processing.

## Code availability

Code required for processing raw reads to OTU tables are available on GitHub: https://github.com/HuttonICS/PineBiomeDataPaper. Code required for calculating the proportion of reads assigned to needle pathogens from the OTU tables is available on GitHub https://github.com/HuttonICS/DNB_Survey_Paper_OTU_processing.

## Author contributions

For the DNB surveys: AP designed, conducted and processed survey results. For the molecular validation: SK, BC and AG created metabarcoding libraries; BM and SJ processed metabarcoding data. For the multisite trial: AP, SC, JB, JS and JC contributed to the multi-site common garden design and management. AP, SC and BM wrote the manuscript and all co-authors contributed to the final version.

## Acknowledgements

We thank Glenn Iason for the original pine trial establishment concept, planning and establishment; Carolyn Riddell, Pete Hedley and Jenny Morris for metabarcoding-related lab work and discussions; Kevin Donnelly, Juan Pablo Lobo-Guerrero Villegas, Krisztian Nemeth, Hattie Barber, Luisa Dickenmann, Martin Mullett and Richard Baden for fieldwork.

